# Accurate and Robust Characterization of Structural Variants at Low Coverages with ARCLID

**DOI:** 10.1101/2025.10.10.681591

**Authors:** Sajad Tavakoli, Rasmus John Normand Frandsen, Sina Majidian, Marjan Mansourvar

**Affiliations:** Department of Biotechnology and Biomedicine (DTU Bioengineering), Technical University of Denmark, Søltofts Plads, Kongens Lyngby, 2800, Denmark; Department of Computer Science and Engineering, Chalmers University of Technology and University of Gothenburg, Gothenburg, Sweden; Science for Life Laboratory (SciLifeLab), Gothenburg, Sweden; Programming, Logic and Intelligent Systems (PLIS), Department of People and Technology, Roskilde University, Roskilde, Denmark

**Keywords:** Structural Variant Calling, Sequencing Coverage, Low Coverage, Long Reads, Deep Learning, YOLO

## Abstract

Structural variants (SVs) shape genome diversity and underlie human diseases, yet accurate detection remains difficult, especially at low sequencing coverages. Most existing callers are optimized for deep sequencing and lose sensitivity as coverage falls, limiting their use in studies that sequence genomes at shallow depth. We present ARCLID, a deep learning-based SV caller that reframes SV detection as an object detection task. We benchmarked ARCLID against six well-established callers on four samples and challenging medically relevant genes, across coverages from 28× down to 5×, multiple SV sizes, and both relaxed and strict breakpoint matching. At 5×, ARCLID achieved the highest F1 score of all evaluated callers on every sample, exceeding the second-best tool by up to 8%. This advantage widened under strict breakpoint matching, indicating genuine variant recovery with accurate breakpoint placement. At higher coverages, ARCLID remains competitive with the strongest current tools. By preserving accuracy at low depth, it makes reliable SV discovery feasible from shallow data, enabling lower sequencing cost, time, and storage for population-scale and clinical genomics. We additionally curated SV benchmarking regions for HG00733 and NA19240 that complement the widely used HG002 GIAB benchmark regions, supporting evaluation of SV callers across ancestrally distinct genomes.

## Introduction

Structural variants (SVs) are large-scale genomic alterations (>50 bp) in the human genome that can significantly influence gene regulation, function, and ultimately phenotypic diversity [1], [2]. These variants are also increasingly recognized as key contributors to human disease, with SVs having been linked to conditions such as Huntington’s disease, driven by pathogenic CAG repeat expansions in the HTT gene [3], and 22q11.2 deletion syndrome (DiGeorge syndrome), caused by recurrent deletions on chromosome 22 [4]. Among SVs, long insertions and deletions are the most frequent forms and have been implicated in diverse genetic disorders [5] and evolutionary genomics [6].

Detection of SVs from genomic sequencing data has traditionally followed two main directions: assembly-based and read alignment-based approaches [7]. Assembly-based methods reconstruct long contiguous sequences (contigs) from raw reads and then compare the assembled genome to a reference to identify SVs. This strategy can better capture complex regions like MHC, enabling precise detection of large and complex SVs, especially in repetitive or challenging genomic regions [7]. However, assembly-based approaches require substantial computational resources and depend on high sequencing coverage and multiple sequencing technologies [7]. In contrast, read alignment-based SV calling detects SVs from sequencing reads mapped to a reference genome, making it more practical for routine applications and lower coverage data [7].

Alignment-based SV calling, especially from long reads, typically follows a multi-step process. First, sequencing reads are aligned to a reference genome using long-read mappers such as minimap2 [8] or pbmm2, which provide detailed information on mismatches, gaps, and breakpoints. From these alignments, characteristic signals are extracted, including split reads [9], discordant read pairs [10], local shifts in read depth[11], and large gaps in CIGAR strings [12]. These raw signals are usually aggregated through clustering algorithms, grouping reads that support the same putative variant and at the same time filtering out noise from sequencing errors. The clustered evidence is refined to improve breakpoint resolution, often by reanalyzing supporting reads or constructing consensus sequences. Candidate variants are subsequently classified into standard SV types. Finally, genotypes are assigned by comparing the proportion of reads supporting the variant versus the reference allele. This traditional approach, employed by tools such as Sniffles [13], Sniffles2 [14], cuteSV [15], and SVIM [16], pbsv [17], and DeBreak [18], is highly dependent on clustering thresholds and hand-crafted features/rules for integrating evidence.

These SV callers detect structural variants by searching for intra-alignment and inter-alignment signatures in aligned reads. Intra-alignment signatures correspond to gaps in either the read or the reference, as indicated by the CIGAR string, while inter-alignment signatures reflect discordances between segments of a split read [16]. In these tools, feature extraction is generally static and platform-specific, requiring manual parameter tuning to accommodate differences in sequencing technologies (e.g., Nanopore vs. PacBio) or sequencing coverage. Furthermore, multiple evidence sources, such as split-read patterns, coverage depth, etc., are often integrated in a sequential or additive manner, potentially limiting flexibility in the analysis. Sniffles2, the latest version of Sniffles, builds on these traditional steps but incorporates a novel algorithm designed to automatically adjust parameters, thereby alleviating the need for manual tuning [14]. Although these well-established SV callers generally provide reasonable accuracy, they often exhibit reduced performance under low sequencing coverage and when detecting very large SVs, primarily due to insufficient read support and alignment ambiguity across extensive genomic rearrangements.

These limitations are particularly important as large-scale genome sequencing initiatives increasingly require accurate variant detection at reduced sequencing depths, where cost, instrument throughput, and data-storage burden become major constraints. Large-scale genome sequencing programs are expanding worldwide. For example, the All of Us Research Program aims to sequence more than one million human genomes and has already generated PacBio HiFi data for over 1,000 samples at approximately 7× coverage [19], [20]. Denmark has also committed about one billion Danish kroner to sequence 60,000 genomes at 30× coverage [21], with similar initiatives emerging in countries such as China [22], Russia [23], and South Africa [24].

In efforts of this scale, reducing coverage from typical ∼30× to ∼5× lowers per-genome sequencing cost by roughly three-to five-fold. Also, it can increase instrument throughput by about four-fold (a single PacBio system can sequence ∼2,500 human genomes per year at 20× coverage versus ∼10,000 per year at 5× — https://www.pacb.com/revio/, accessed May 2026), and shrink storage requirements from ∼100 GB to ∼15 GB per genome at HiFi quality. This throughput gain follows directly from how sequencing capacity is allocated. Because the output of a PacBio SMRT Cell is finite, sequencing a single human genome to ∼30× coverage may consume close to the full output of one SMRT Cell, depending on sequencing chemistry and run yield. When lower coverage, such as 5×, is sufficient, each sample requires only a small fraction of the total data output. Therefore, barcoding and multiplexing allow several genomes to be pooled on the same SMRT Cell, improving sample throughput and substantially reducing the sequencing cost per genome. Consequently, a computational method that preserves variant calling accuracy at substantially lower coverage could deliver disproportionate benefits across population genomics, clinical diagnostic laboratories, academic research groups, newborn-screening programs, and cancer centers.

In this study, we introduce ARCLID to address the mentioned gaps. Our tool is designed for the low-coverage PacBio HiFi sequencing, where existing callers degrade most sharply. As sequencing depth decreases, ARCLID sustains detection sensitivity that competing tools lose. At 5× coverage, it achieves the highest F1 score of all evaluated callers on every sample tested, improving accuracy up to 8%. Critically, this advantage holds under both relaxed and stringent breakpoint-matching criteria, which indicates that the gain reflects genuine variant recovery rather than lenient evaluation. At higher coverages, where SV detection is already well served by existing methods, ARCLID performs on par with the strongest current tools. The practical implication is direct. Because ARCLID preserves calling accuracy at depths as low as 5×, it enables reliable SV detection from data that would otherwise be considered too shallow, which substantially reduces the sequencing cost, instrument time, and storage burden of large-scale and clinical genomic studies.

The remainder of the manuscript is organized as follows. In the Materials and Methods section, we describe the design of ARCLID and the implementation of each component of the pipeline. We then comprehensively benchmark ARCLID against several well-established SV callers across a diverse set of samples, sequencing coverages, and SV size categories, using two breakpoint evaluation strategies (relaxed and strict matching criteria). The corresponding evaluation results are presented in the Results section.

## Materials and Methods

### Overview

ARCLID is an end-to-end framework that reformulates structural variant calling from long-read alignments as an object detection problem in computer vision. DeepVariant [25] previously demonstrated that convolutional neural networks can learn patterns associated with SNPs and small indels from images constructed using aligned-read pileups. Building on the broader concept of image-based variant calling, we extend this paradigm to long-read SV calling and formulate their detection as an object-detection task. Our innovation lies in how we construct the image; ARCLID restructures complementary alignment signals and encodes them as three-channel pileup images, enabling object detection–based deep learning models to learn complex patterns associated with SV breakpoints, variant types, and genotypes directly from the data. The ARCLID workflow (Figure 1) consists of four main steps: (1) extracting alignment features, (2) generating three-channel pileup images, (3) localizing, classifying, and genotyping SVs using two YOLOv11x models [26], and (4) postprocessing the resulting predictions. Each step is described in detail in the following subsections.

**Figure 1.**
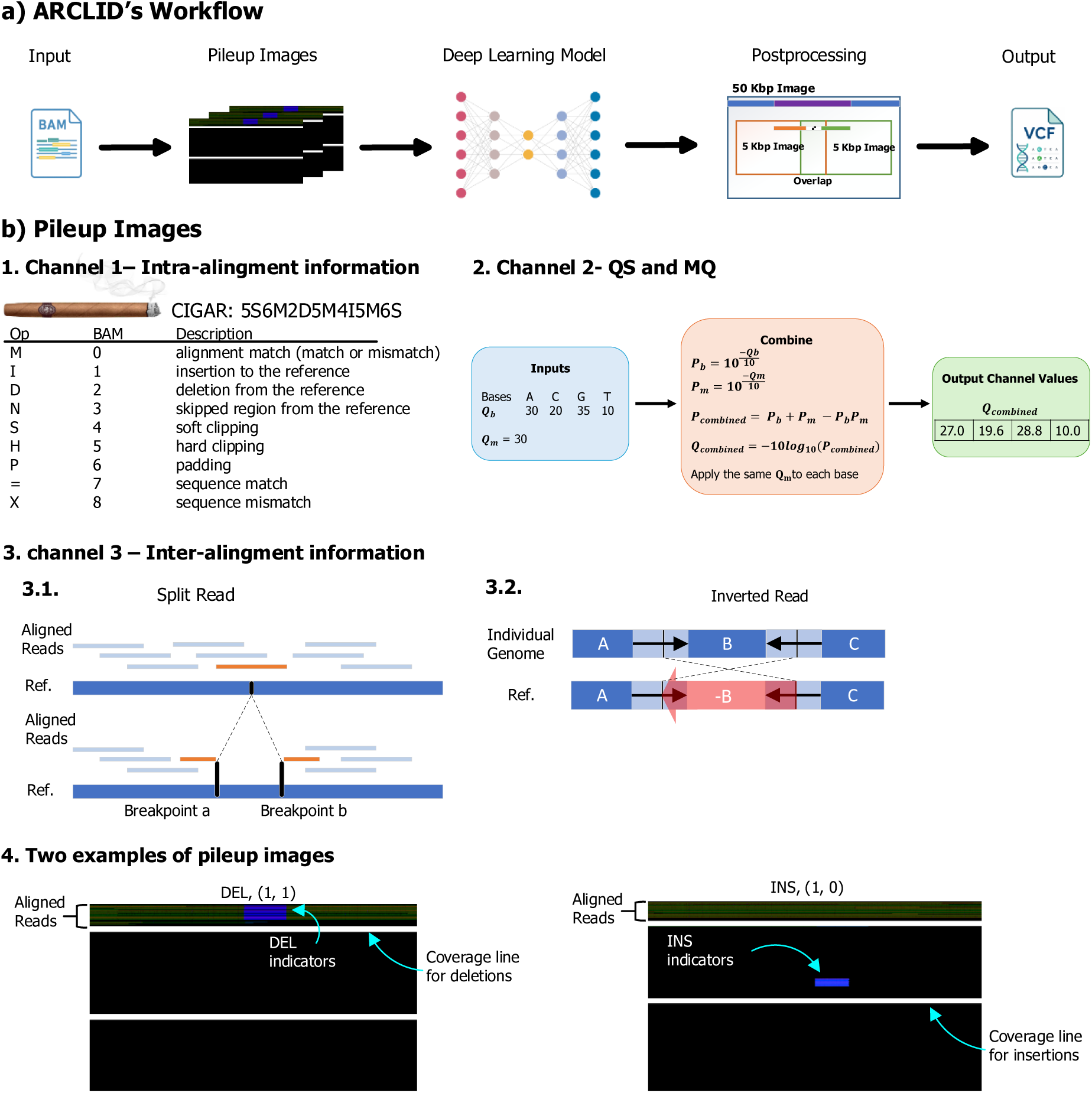
The Overview of the pileup-image representation and ARCLID workflow. (a) ARCLID workflow. Pileup images are generated from read alignments, analyzed by a deep learning model, and then post-processed across overlapping windows to produce the final VCF output. (b) Pileup-image design used to encode read-alignment evidence for structural variant calling. Channel 1 represents intra-alignment information derived from CIGAR operations, including matches, insertions, deletions, clipping, and related alignment states. Channel 2 combines base-quality scores and mapping-quality scores into a single quality-informed channel. Channel 3 encodes inter-alignment information, including split-read signals and inverted-read patterns. Representative deletion and insertion pileup images illustrate how aligned reads, variant indicators, and coverage lines are represented.

### Feature Extraction and Pileup Images

Two sets of pileup images are generated. The first set covers regions of 5 Kbp with a 1 Kbp overlap, designed to capture shorter indels (<2 Kbp). The second set spans 50 Kbp with a 10 Kbp overlap, intended to capture longer indels (>2 Kbp). The 5 Kbp images are used as input to the first YOLO model, while the 50 Kbp images are used for the second model. We generate two sets because very long SVs can span several pileup images, making them difficult to detect once divided.

Each image is constructed to support a sequencing depth of up to 150 reads per position. In these images, columns correspond to each base pair position in the reference genome, and rows correspond to aligned reads. The first channel stores insertions, deletions, and soft-clipped flags, the second channel is dedicated to the combination of quality scores and mapping qualities, and the third channel is to highlight inter-alignment signals (if a read is split, supplementary, or reversed).

#### The first channel of pileup images

The first channel of the pileup images stores intra-alignment information presented in CIGAR string such as “D”, “I”, and Soft Clip flags (“S”). Generating the pileup image is not straightforward because deletions and insertions introduce positional shifts between the reads and the reference sequence. Therefore, simply loading the aligned reads into the pileup image from the first to the last aligned position does not preserve the correct genomic coordinates. This can result in a discrepancy between the displayed SV position and its corresponding label in the ground-truth VCF file. Therefore, careful handling of this process is essential to prevent misalignment, which could mislead the deep learning model during training. To address this issue, deletion flags (’D’) are incorporated alongside aligned sequences in columns 0–150 with the intensity of 200, while base pairs associated with insertion flags (’I’) are extracted from the reads and placed in columns 150–300 with the intensity of 250. This approach effectively mitigates positional shifts introduced by insertions, ensuring accurate SV representation in the pileup images.

Finally, a horizontal line is drawn to represent the average coverage of the sample. This coverage line is drawn from the row equivalent to the average coverage of the sample and helps the deep learning model to better understand coverage variations. Examples of the generated pileup images are presented in Figure 1.b.4.

#### The second channel of pileup images

To combine mapping quality and quality scores for individual base reads, the two scores (from the aligner and sequencing machine, respectively) are converted from PHRED score into probabilities of error (Equations 1, and 2).

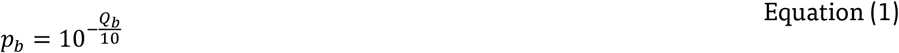

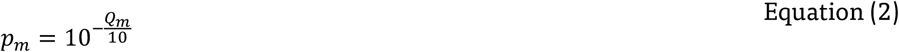

where a base call has a PHRED quality score *Q_b_*, it represents the probability of base-call error *p_b_*. Similarly, the mapping quality *Q_m_* describes the likelihood that the read is incorrectly mapped. Its associated probability of mapping error *p_m_*.

There are now two probabilities of error: one from the base call and one from the mapping. Assuming (for simplicity) that these errors are independent events, we can combine them to find the overall error probability for a given base alignment. The error could occur if either: a) The base is called incorrectly, b) The read is mapped incorrectly, or c) Both. Under independence, the combined error probability *p_combined_* can be expressed using the complement rule:

- Probability that the base is correct: 1 − *p_b_*
- Probability that the read is mapped correctly: 1 − *p_m_*

Thus, the probability that either is wrong (the combined error probability) is computed by Equation (3), and finally, we convert it back into PHRED score through Equation (4).

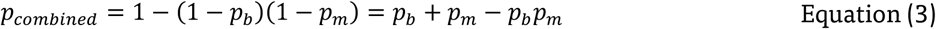

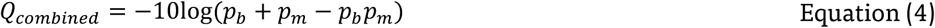

#### The Third Channel of Pileup Images

The third channel is to highlight inter-alignment signatures. To encode these signatures, we used a decision tree to classify each read. Figure 2 shows the binary decision tree designed to encode reads based on whether they are split, supplementary, or mapped in reverse orientation. Figure 2 illustrates the decision tree, including the root node (“is split?”), the internal nodes (“is supplementary?” and “is reversed?”), and the leaf nodes (“111”, “110”, etc.). In this decision tree, certain states are logically impossible, for example, cases where a read is not split but is marked as supplementary. Clearly, a read cannot be supplementary if it is not split. Therefore, the two states 011 and 010 cannot occur. Leaf nodes demonstrate how a binary value is assigned to a read, then the binary value is converted to decimal one, and ultimately the value is multiplied by 20 to increase the separation between values, thereby helping the deep learning model better differentiate them.

**Figure 2.**
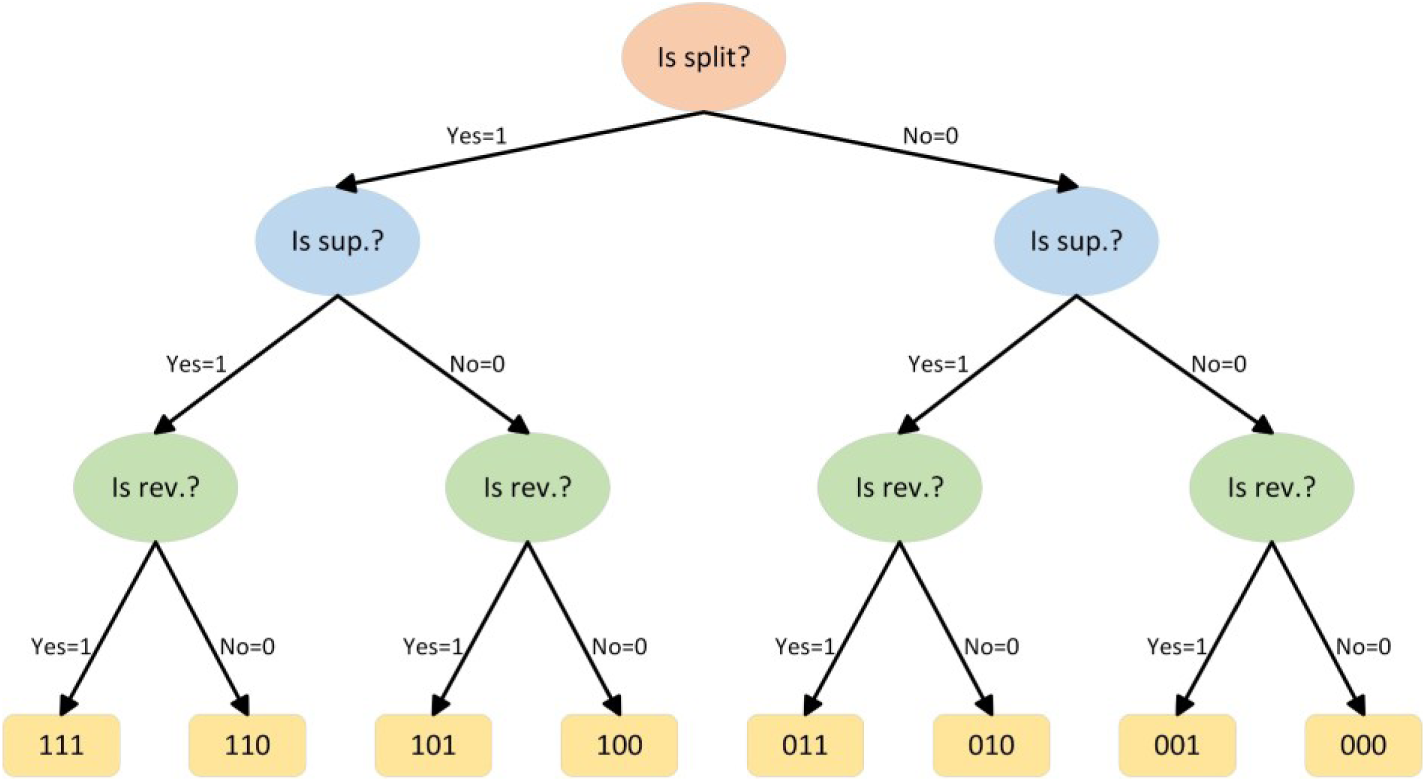
Decision tree for categorizing reads as split, supplementary, or reversed. The decision tree begins at the root node, which determines whether the read is split. The subsequent internal nodes check whether the read is supplementary or reversed. At the leaf nodes, each read is assigned a binary value. Finally, these binary values are converted to decimal form and multiplied by 20 to enhance the distance between categories.

### Datasets and Evaluation Strategy

We benchmarked ARCLID alongside well-established SV callers, Sniffles2 [14], CuteSV [15], SVIM [16], pbsv [17], DeBreak [18], and SVision-pro [27]. We used PacBio HiFi data of HG002 (released by GIAB) [28], HG00733 and NA19240 (from HGSVC) [29], simulated data generated by VISOR [30], as well as targeted analysis on challenging medically relevant genes (CMRG) from GIAB [31]. We chose the best and the most well-known SV callers published in top-tier journals for performance comparison. Among the six competitors, Sniffles2, CuteSV, SVIM, pbsv, and SVision-pro are alignment-based SV callers, whereas DeBreak combines alignment information with local de novo assembly. Since ARCLID is also based on deep learning for object detection, we included SVision-pro in the comparison to evaluate ARCLID against deep learning-based SV caller. We also attempted to include SVHunter [32], another deep learning-based SV caller, and volcanoSV [33], an assembly-based SV caller. However, both tools were extremely difficult to install and did not produce usable results even after installation. For example, SVHunter generated around 300 GB of intermediate data while processing a 5× coverage sample. We also encountered many errors and bugs when running volcanoSV during contig loading, and although we made substantial efforts to fix the issues and rebuild the tool, we were ultimately unable to run it successfully.

We adopted a chromosome-based data partitioning strategy to separate the training and test datasets. Chromosomes 1-10 of HG002 were used to train the models, chromosome 11 for internal evaluation, and chromosomes 12-22 for benchmarking. Sample HG002 from GIAB Tier 1 provides high-confidence regions and includes ∼12,000 long insertions and deletions, making it a reliable standard for evaluating variant calling performance [28]. Furthermore, simulated data containing synthetic variants provides a reliable resource for benchmarking, as these datasets are generated based on pre-defined variant sets, where the exact type and location of each variant are already known. As a result, the ground truth is unambiguous, allowing for precise evaluation of variant calling methods without the uncertainty that may be present in real sequencing data. The entire process to generate simulated PacBio reads is provided in Supplementary Information S3.

To expand our benchmarking, we also included two HGSVC samples, namely, HG00733 and NA19240, with over 25,000 SVs per sample, making rich truth sets for SV discovery. However, these samples lack validated benchmarking regions that could avoid problematic regions and enhance the reliability of the benchmarking (similar to high-confidence regions that is available for HG002 provided by GIAB [28], [29]). This is particularly important because the measured performance and relative ranking of SV callers can vary substantially depending on the selected ground-truth callset, reference genome, and genomic regions included in the evaluation [34]. To make these rich truth sets usable for a fair benchmarking for all tools, we curated SV regions for HG00733 and NA19240 (detailed in the next subsection).

### Curation of HGSVC benchmarking regions

Inspection of the assembly-based VCFs revealed loci in which two or more SVs of significantly differing size or type were packed within one to two kilobases, sometimes accompanied by partial or absent genotypes. At such complex loci, differences in variant representation can confound variant-level benchmarking. For example, a caller may represent a complex event as a single SV, whereas the truth set may encode it as several nearby records. Because Truvari [35] (a well-known tool for benchmarking SV callers) generally performs one-to-one matching, only one truth record may be matched, while the remaining records are counted as false negatives. Also, if the caller’s representation is only matched with a missing genotype record, it may additionally be counted as a false positive because missing genotype records are filtered out. Consequently, errors at these loci may partly reflect representation ambiguity rather than genuine detection failure. Thus, in these regions, a caller that correctly reports one variant but not its close neighbors is penalized by Truvari [35] with spurious false negatives or false positives that reflect annotation density rather than caller performance.

The clustering of variants observed in these truth sets is an expected property of assembly-based callsets rather than an indication of erroneous calls. SVs are strongly enriched in tandem repeats and segmental duplications, where alignment ambiguity permits several equivalent representations of the same underlying event [35], [36]. Alignment of a diverged segment frequently yields a deletion of reference sequence followed immediately by an insertion of assembly sequence, and phased callsets can emit near-identical variants on each haplotype when breakpoint placement differs slightly between them [37].

To address this, we define low-ambiguous SV regions for each sample as follows: we applied single-linkage clustering to the variant footprints on each chromosome and discarded any cluster containing two or more variants, retaining only well-isolated loci. We additionally excluded sites carrying missing or partially missing genotypes. The surviving isolated SV regions, expanded by a small flank, were written to a BED file and fed to Truvari [35] through its *includebed* parameter so that performance is measured only where a one-to-one correspondence between the truth and the call can be established unambiguously. Because the curation depends only on the published VCFs by HGSVC [29] and a fixed distance threshold, the resulting region sets are deterministic and can be used by others. We therefore release the cleaned HG00733 and NA19240 benchmarking regions as a valuable resource that complements the widely used HG002 GIAB region. This extends rigorous, low-ambiguity structural variant evaluation to two additional, ancestrally distinct genomes.

### Deep Learning Model and Training

The deep learning model employed in this study is an object detection framework commonly used in computer vision tasks. We used the YOLOv11x architecture [26], which comprises 56.9 million parameters, and trained it using the HG002 sample from the GIAB consortium due to the high accuracy and reliability of its labels. Chromosomes 1–10 were allocated to the training set, chromosome 11 to the evaluation set, and chromosomes 12–22 to the testing set. HG002 HiFi data with various coverage levels (28×, 20×, 10×, 5×) were employed for training. It is worth noting that the other samples used in this study—HG00733, NA19240, and the simulated sample—were excluded from training and treated as independent test samples to assess the **generalizability** of ARCLID.

The models were trained for 200 epochs with a batch size of 16, using pre-trained weights. During the initial training phase, the models failed to converge. By comparing the pileup images with the truth-set coordinates, we found that about 30% of SVs were shifted from their annotated positions, in some cases by more than 1 kbp. This mismatch likely reflects the fact that the GIAB truth set was generated using a combination of sequencing platforms, aligners, SV callers, and phasing pipelines, whereas our pileup images were produced from a single sequencings platform aligned with a single mapper. To address this issue, we programmatically drew ground-truth bounding boxes on all SVs and manually curated approximately 50,000 images that took us around three months. We then retained for training only the images in which the SV signal was located within the annotated bounding box. Retraining the models on this curated dataset led to proper convergence and substantially improved validation and test performance (Supplementary Information Supplementary Figure 1).

### Postprocessing

Following the processing of YOLO’s outputs, variants located within 100 base pairs and sharing the same SV class and genotype are merged. Subsequently, the candidate variants undergo a refinement process by revisiting their genomic positions within the pileup image. During this step, the variants are evaluated based on the existing indicators of ‘ D ‘ and ‘ I ‘ signals from CIGAR string present in the image. A variant is retained only if these indicators align with the predictions provided by the model. Finally, the resulting variant information is systematically organized and recorded in a Variant Call Format (VCF) file.

### Data Preparation

The alignment BAM file for HG002 (aligned to the hg37 reference genome using pbmm2) is publicly available; therefore, we directly employed this file without realigning the raw reads. We downloaded the raw reads for NA19240 and HG00733 and aligned them to the hg38 reference genome using pbmm2 v 1.17.0. Simulated reads were generated from the hg38 reference genome using VISOR. Detailed procedures for read mapping, generating simulated data, the command line to run all SV callers are provided in the Supplementary Information.

### Benchmarking Modes and Metrics

To benchmark ARCLID and other available SV callers, we employed Truvari [35] in two evaluation modes. The stringency of variant matching in Truvari can be controlled through three key parameters: *refdist*, *pctsize*, and *pctovl*. The *refdist* parameter defines the maximum allowed distance (in base pairs) between the breakpoints of real SV and detected SV to be considered a match. The *pctsize* parameter specifies the minimum required similarity in variant size (expressed as a percentage), and *pctovl* determines the minimum reciprocal overlap between real SV and detected SV. Together, these parameters control the strictness of the comparison and enable benchmarking under both relaxed and strict evaluation conditions.

In the strict mode, we required highly stringent matching criteria with parameters set to *refdist* = 50 *bp*, *pctsim* = 0.9, and *pctovl* = 0.9, ensuring precise breakpoint agreement and allele similarity between predicted and reference variants. In the relaxed mode, we applied more tolerant criteria with *refdist* = 1 *Kbp*, *pctsim* = 0.7, and *pctovl* = 0, allowing for approximate matches and broader overlap tolerance. This dual approach enabled us to assess performance under exact-match conditions as well as under more flexible benchmarking conditions typically used in other studies, where differences in breakpoints or allele similarity are still considered acceptable. The full results for both benchmarking modes across all coverages and SV sizes are available in Supplementary Tables 1-20.

We used four metrics to capture/describe the tools’ performance which are precision (Equation 5), recall (Equation 6), F1-score (Equation 7), and true positive genotyping rate (TP-GT Rate, Equation 8). The TP-GT rate represents the proportion of SVs that are not only correctly detected and classified but also correctly genotyped, relative to the total number of SVs.

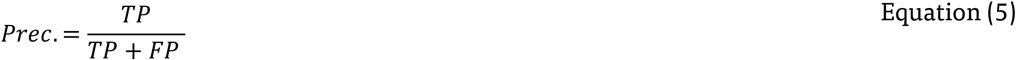

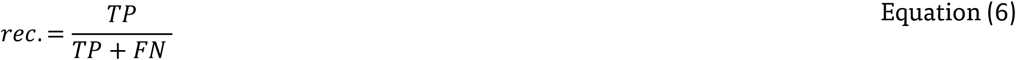

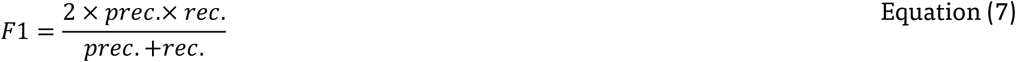

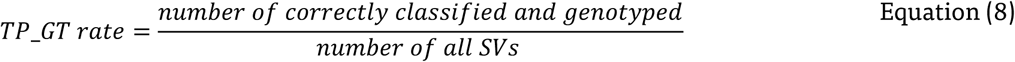

In the equations 5-8, *TP* (true positives) denotes the number of SVs correctly identified by the method, *FP* (false positives) represents the number of incorrectly predicted SVs that are not present in the ground truth, and *FN* (false negatives) refers to the number of SVs present in the ground truth but missed by the method.

## Results

### ARCLID outperforms state-of-the-art methods for SV calling

We evaluated ARCLID against six widely used structural variant callers, namely Sniffles2 [14], cuteSV [15], SVIM [16], pbsv [17], SVision-pro [27], and DeBreak [18]. The evaluation included three real samples, HG002, HG00733, and NA19240, as well as one simulated sample. We also assessed performance using the GIAB Challenging Medically Relevant Genes (CMRG) benchmark, which contains curated, difficult, and medically important loci in HG002. All tools were evaluated across a range of sequencing coverages and under both relaxed and strict breakpoint matching criteria. Figure 3 provides a schematic overview of the evaluation, while Supplementary Tables 1–20 present the numerical results.

**Figure 3.**
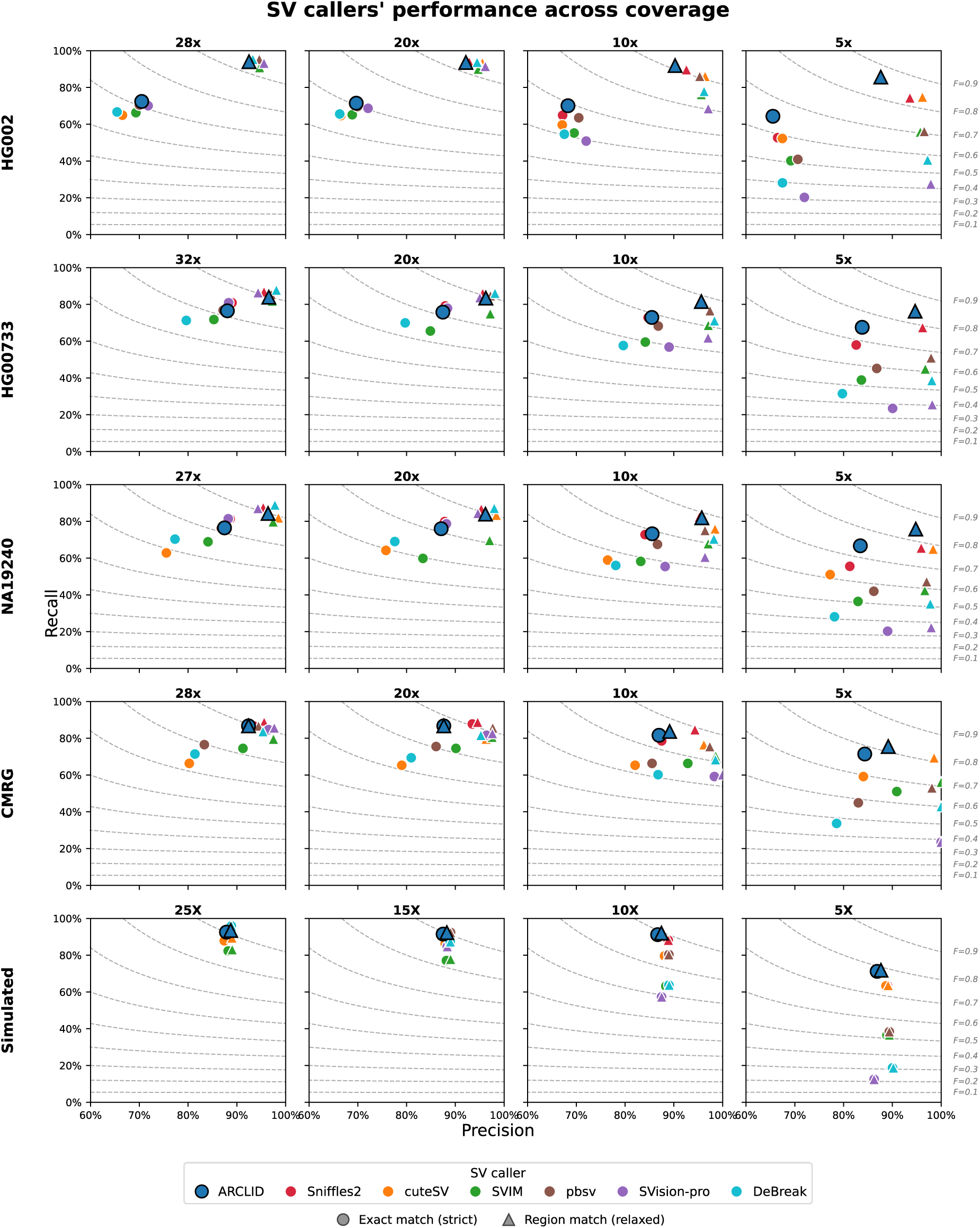
Precision-recall performance of SV callers across coverage under strict and relaxed matching. Recall versus precision for ARCLID (blue) and six other SV callers, by evaluation set (rows: HG002, HG00733, NA19240, GIAB CMRG, and simulated data) and sequencing coverage (columns, decreasing left to right). Circles show exact matching (strict) and triangles show region matching (relaxed), two strategies to evaluate breakpoints. Dashed curves are iso-F1 contours (0.1–0.9). ARCLID sits in the high-precision, high-recall region across samples, with a growing advantage at low coverage (10× and 5×).

Across all evaluation sets, ARCLID exhibited consistent highly accurate performance in which its relative advantage grew as coverage decreased. At high coverage, where SV detection is already well resolved, ARCLID performed on par with the strongest available methods, and the leading callers clustered within a narrow band of F1 scores. On HG00733 at 32×, in strict mode evaluation, the top methods fell within an F1 range of approximately 82% to 85%, and on the CMRG benchmark at 28× the three leading callers were separated by less than 1% F1, indicating that broadly similar performance at high coverages. As coverage was reduced to 10× and 5×, ARCLID achieved the highest F1 score of all evaluated callers across every evaluation set. At 5×, ARCLID exceeded the second-best method by approximately 5% F1 on HG002, 7% F1 on HG00733, and 8% F1 on NA19240, showing that its accuracy degrades more gracefully than that of existing tools as read support becomes limited.

The same coverage-dependent pattern held on the CMRG benchmark, where the leading methods were closely grouped at high coverages (28× and 20×), while ARCLID achieved the highest F1 at 10× and 5×, indicating reliable structural variant recovery in medically relevant genes even at low sequencing depth. This trend was further reproduced on simulated data, where ARCLID performed comparably to the most accurate methods at high coverage and attained the highest F1 at 10× and 5×, supporting the conclusion that the low-coverage advantage of ARCLID is a robust property of the method rather than a feature of any single dataset.

### ARCLID Accurately Localizes SV Breakpoints

To distinguish accurate variant detection from accurate variant localization, we evaluated every caller under two matching criteria for breakpoint localization. Relaxed matching credits a call when a variant of the correct type is reported in the approximate vicinity of a true event, and therefore primarily measures detection. Strict matching additionally requires close agreement between the reported and reference breakpoints, and therefore jointly measures detection and breakpoint localization. Strict matching is especially critical for clinically and functionally important genomic regions, where even small positional errors can alter biological interpretation [38]. For instance, in oncogenomics, the exact localization of breakpoints is essential for identifying gene fusions (e.g., BCR–ABL1, TMPRSS2–ERG) [39], [40], which drive cancer progression and guide targeted therapy decisions. A method that reports the presence of a variant but misplaces its boundaries provides limited value in these settings.

ARCLID performed strongly under this stricter criterion. Its advantage over competing methods at low coverage was larger under strict breakpoint matching than under relaxed breakpoint matching across all samples. As shown in Figure 4, at 5×, the F1 advantage of ARCLID over the second-best caller increased from approximately 2.5% to 6% on HG002, from approximately 5.2% to 6.7% on HG00733, and from approximately 6% to 8.2% on NA19240 when moving from relaxed to strict breakpoint matching. The widening of this margin under more stringent breakpoint requirements indicates that part of the ARCLID advantage derives from more accurate breakpoint placement rather than from detection alone.

**Figure 4.**
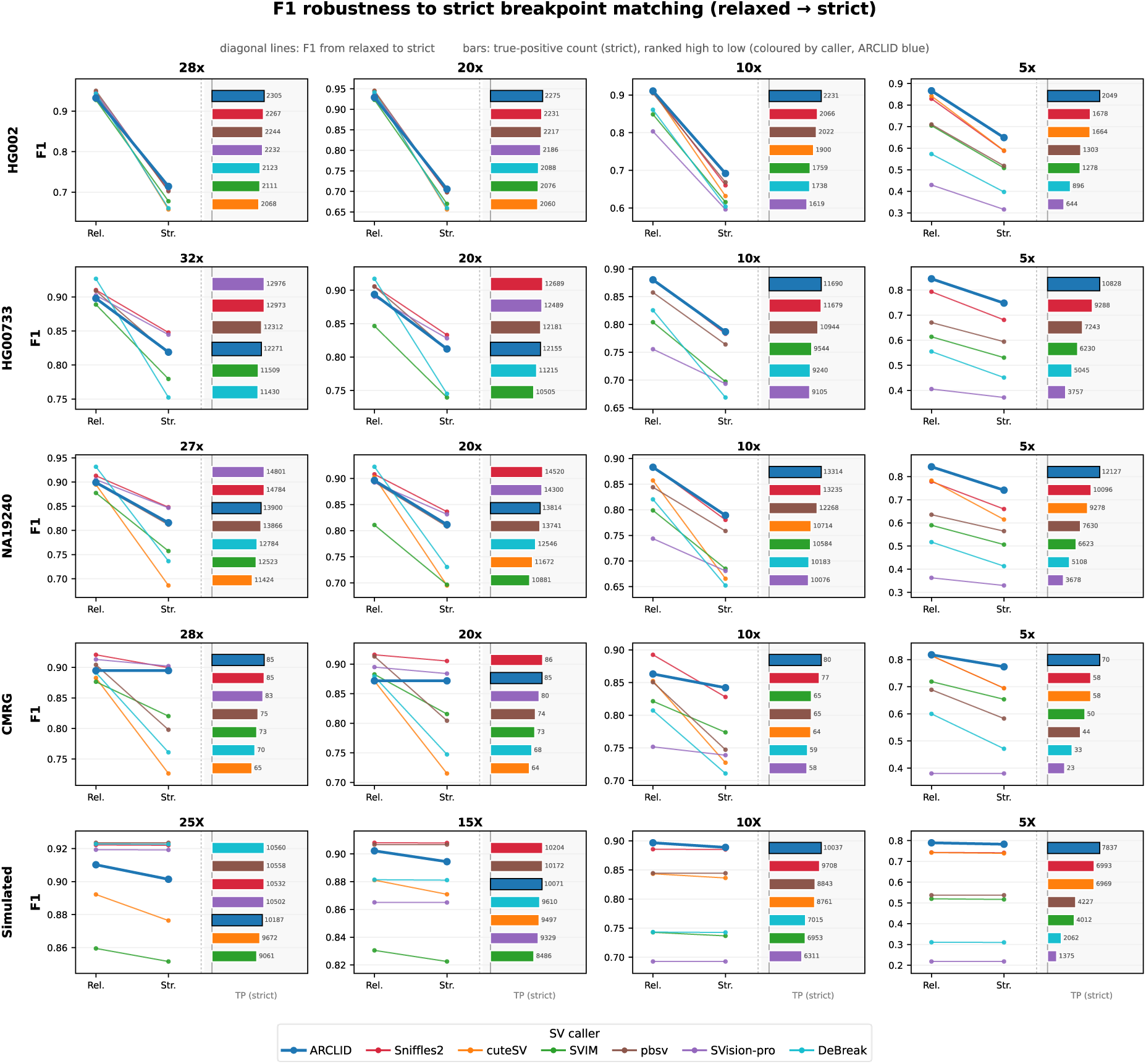
F1 robustness to strict matching and true-positive yield across coverage. For each caller, diagonal lines show F1 from relaxed (“Rel.”) to strict (“Str.”) matching, and horizontal bars show the strict-mode true-positive count (coloured by caller, ranked high to low, ARCLID in blue). Rows are evaluation sets (HG002, HG00733, NA19240, GIAB CMRG on HG002, simulated data). At high coverages (>=20×), ARCLID’s performance is on par with other callers, while at 10× and 5×, ARCLID retains high strict-mode F1 and high true-positive yield across samples.

We further quantified breakpoint localization directly as the proportion of detected variants whose breakpoints agreed closely enough with the reference to satisfy strict matching, computed as the ratio of recall under strict matching to recall under relaxed matching. On HG002, ARCLID retained the highest proportion of its detections of any caller at every coverage, ranging from approximately 77% at 28× to 75% at 5×, demonstrating superior breakpoint accuracy on this sample. On HG00733 and NA19240, ARCLID likewise retained a high proportion of its detections under strict matching, between approximately 88% and 91% across coverages, placing it among the most accurate methods. The few callers that retained a marginally higher proportion achieved this only on substantially smaller and more conservative detection sets. For example, at 5× on HG00733, Svision-pro retained a slightly higher fraction of its calls but recovered only 23% of true variants, whereas ARCLID retained a comparable fraction while recovering 67.5% of true variants. ARCLID therefore combined high breakpoint fidelity with the highest sensitivity among all evaluated methods, a combination that no competing caller achieved at low coverage. Taken together, the strict-mode results indicate that ARCLID not only detects more true variants at low coverage but also localizes their breakpoints with high precision, a feature most relevant to functional interpretation and clinical application.

### ARCLID Combines High Precision with Superior Recall at Low Coverage

To characterize the basis of the low-coverage advantage, we examined precision and recall separately in addition to their harmonic mean, the F1 score. The two contributions differed in their relative importance across samples, but in every case ARCLID achieved the most favourable overall balance. On HG002, ARCLID achieved the highest recall and the highest F1 at every coverage under strict mode evaluation. Precision differences among the evaluated callers were small on this sample, with all methods falling within a narrow band, for example approximately 66% to 72% at 5× (under strict mode evaluation), whereas recall varied widely, from approximately 20% to 64% at the same depth. Because the methods were closely matched in precision, recall was the determining factor in overall accuracy, and the substantial recall advantage of ARCLID translated directly into the best F1. At 5×, ARCLID recovered approximately 12% more of the true variants than the next most sensitive method while maintaining precision comparable to the field, demonstrating that it converts limited read support into reliable detection more effectively than existing tools.

On HG00733 and NA19240, ARCLID combined high precision with the highest recall at low coverage. Under strict breakpoint matching, ARCLID precision remained between 84% and 88% on HG00733 and between 83% and 87% on NA19240 from the highest to the lowest coverage, placing it among the most precise callers at every depth. At 5×, ARCLID attained both the leading recall and the leading F1 while preserving precision within a small margin of the most precise method. The few callers that reached marginally higher precision did so at a substantial cost to sensitivity. On HG00733 at 5×, Svision-pro and pbsv reached precision of 90% and 87% but recovered only 23% and 45% of true variants, whereas ARCLID combined a precision of 84% with a recall of 67%. A similar pattern was observed on NA19240, where the most precise competing method recovered approximately 20% of true variants compared with 67% for ARCLID.

Taken together, these results show that ARCLID occupied the most favourable region of the precision-recall space across all samples. On HG002 it delivered a decisive recall advantage while remaining within the common precision band, and on HG00733 and NA19240 it combined high precision with the highest recall, a balance that no competing method achieved at low coverage. The consistency of this behaviour under both relaxed and strict breakpoint matching indicates that the low-coverage advantage of ARCLID reflects genuine recovery of true variants rather than a relaxation of calling stringency.

### Performance by SV type

We next assessed deletions and insertions separately on HG002 (Figure 5, Supplementary Tables 3-4). For deletions, ARCLID achieved the highest F1 at every coverage, with an advantage that widened as coverage decreased and reached approximately 8% F1 over the second-best method at 5×. For insertions, ARCLID was the leading method at 28×, 10×, and 5×, and at 20× it remained within less than 1% F1 of the top method, indicating effectively equivalent performance at that depth.

**Figure 5.**
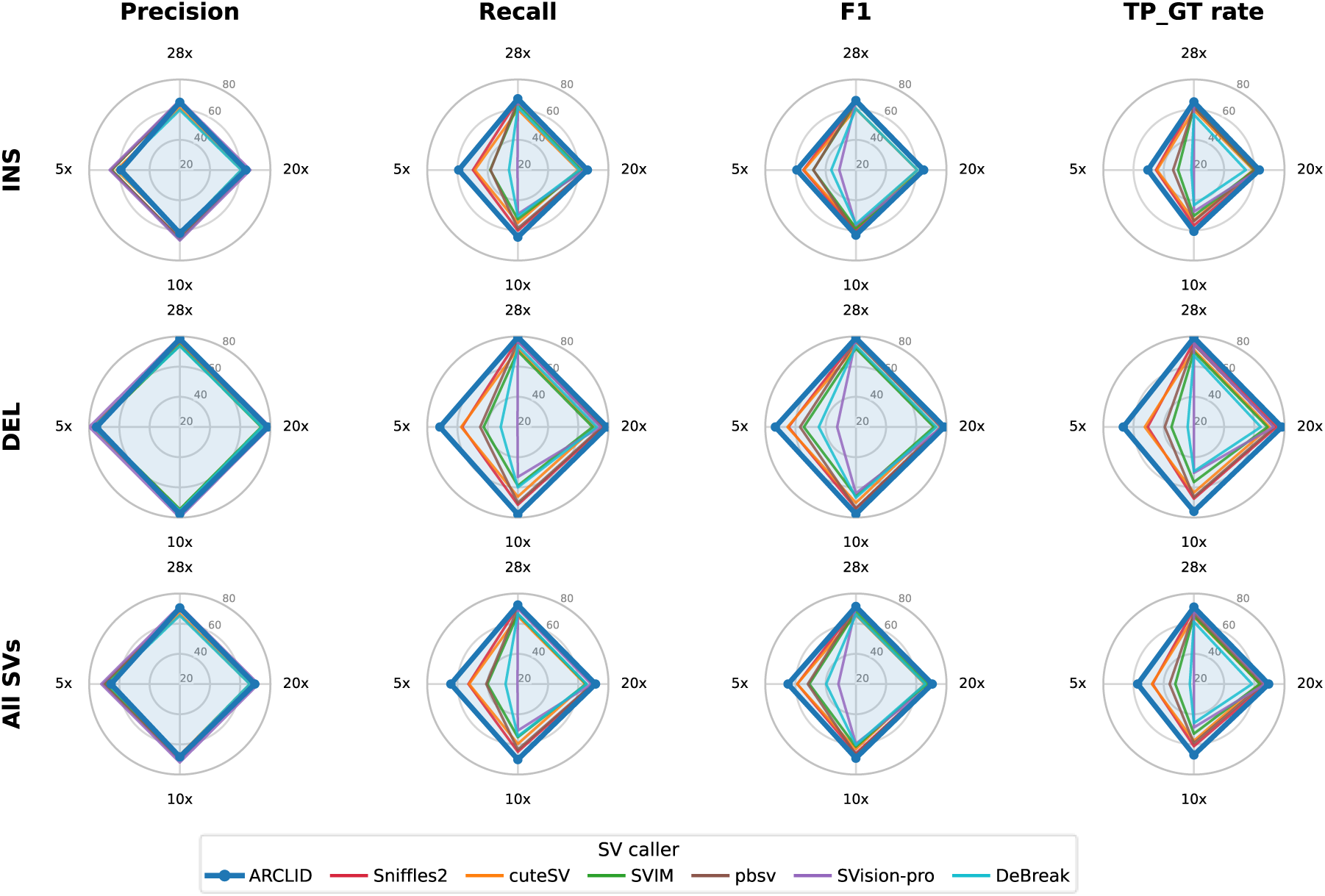
SV callers’ performance on INS and DEL in HG002 under strict matching. Radar plots of precision, recall, F1, and genotype-aware true-positive rate (TP_GT rate) for seven SV callers, by variant type (rows: insertions, deletions, all SVs) and metric (columns). Radar axes are sequencing coverages (28×, 20×, 10×, 5×). ARCLID matches or exceeds competing callers across metrics, most clearly at low coverage and for deletions.

Genotyping accuracy followed the same favourable pattern. For deletions, ARCLID achieved the highest genotype concordance of all evaluated callers at every coverage, remaining above 97% down to 10× and above 93% even at 5×, which indicates that the genotypes assigned to its true detections are highly reliable. For insertions, genotype concordance remained high in absolute terms across the closely grouped field, staying above 85% even at 5×. Because ARCLID combined this accurate genotyping with the highest insertion recall, the overall rate of true insertions that were both detected and correctly genotyped was the highest of all callers at every coverage, and the same held for deletions. ARCLID therefore not only recovers the most true variants at low coverage but also assigns their genotypes correctly more often than competing methods across the genome, which together demonstrate robust performance across the two most common classes of structural variants, with a particularly strong advantage for deletions at low coverage.

**Figure 2.**
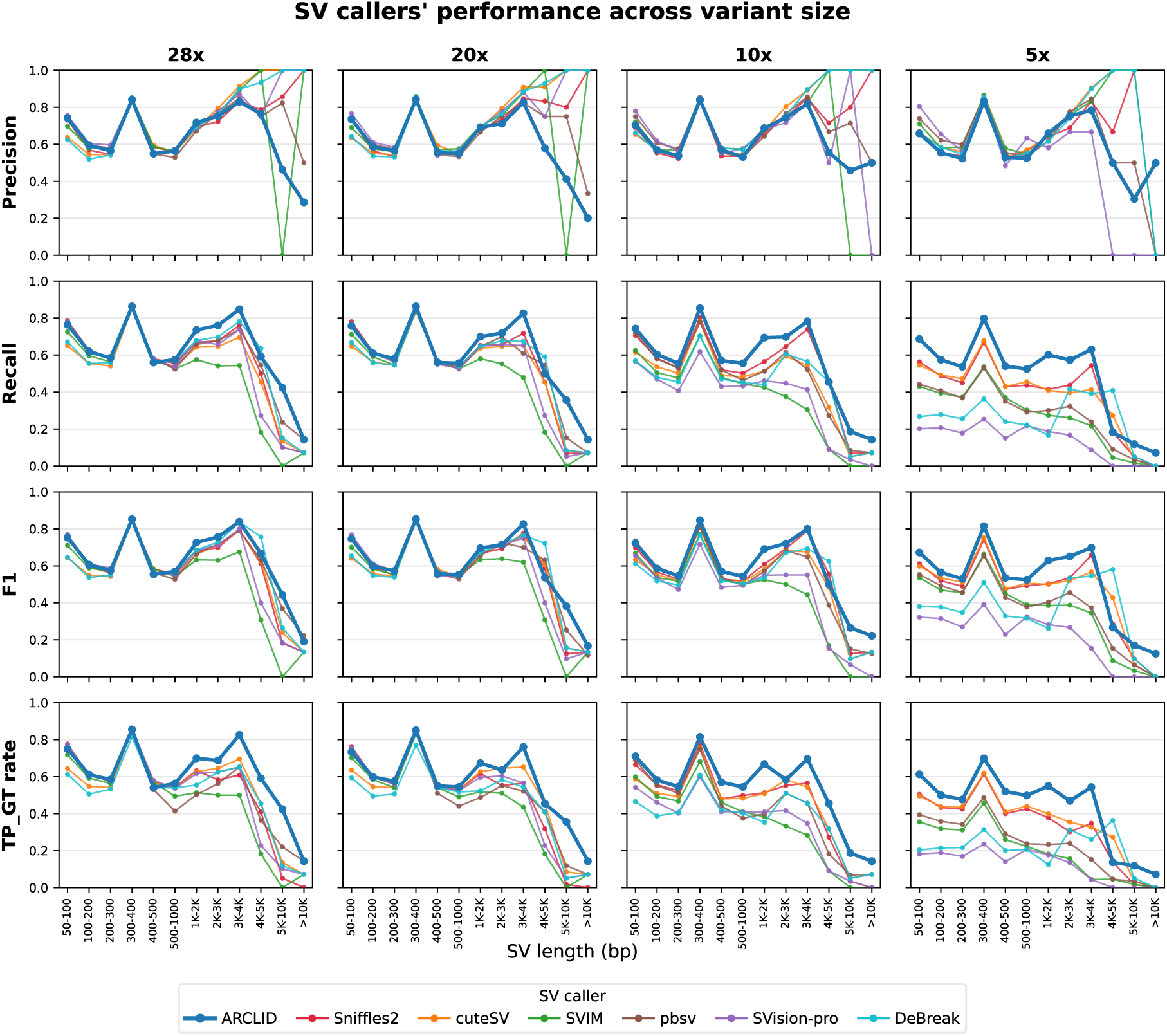
ARCLID’s performance across SV size for all coverages on the HG002 (strict mode) datasets. Columns show sequence coverages (28×, 20×, 10×, 5×); rows show Precision, Recall, F1, and TP-GT rate. ARCLID maintains the highest F1/recall and TP-GT rate across coverages, with the advantage most pronounced for long SVs (≥1 kb) and at low coverage (10×–5×). All tools show declining performance for very large SVs, but ARCLID degrades less severely than other tools. It is also worth noting that the precision of some tools for very large SVs appears to jump to 100% (e.g., for SVs >10 kb, Sniffles2, CuteSV, and SVIM at 28× coverage; and similarly for SVs >4 kb at other coverages). This occurs because these tools detect only a single SV and miss all other SVs. Consequently, according to the definition of precision 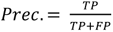, their reported precision becomes very high, in some cases reaching 100%. While this value is technically correct, it is misleading in practice because it does not account for the many undetected SVs, which are reflected instead in the recall.

### Performance Across SV size

To evaluate performance as a function of variant size, we stratified HG002 calls into twelve size classes ranging from 50 to 100 base pairs up to greater than 10 kilobases and repeated the evaluation at each coverage (Figure 6 and Supplementary Tables 13-20). At 5×, ARCLID achieved the highest F1 in nearly all size classes, including the large and challenging categories above 1 kilobase, demonstrating that its low-coverage advantage extends across the full spectrum of variant sizes. At high coverages, ARCLID was the most accurate tool (F1) for medium and large variants and performed comparably to the leading callers for the smallest size classes, where all methods achieved similar accuracy. The retention of accuracy for large variants at low coverage is particularly notable, as these events are typically the most sensitive to reduced read support.

### Impact of Input Features on SV-Calling Performance

To assess the contribution of each design decision, we performed an ablation analysis and evaluated successive versions of the pileup representation on HG002 using chromosomes 12 to 22 at 28× coverage in relaxed mode. The initial representation (version1) encoded deletion and insertion signals in two channels and the combined base-quality and mapping-quality scores in a third channel, while per-position coverage was recorded in the first row rather than as an explicit coverage line. This version achieved a precision of 81% and a recall of 83%, which was reasonable on its own but still below the performance of established callers. Adding an explicit average-coverage line (version 2), which gave the model a direct reference for local depth variation, improved both precision and recall by about 4%. Encoding inter-alignment signatures (version 3) such as split, supplementary, and reverse-orientation states through the decision-tree scheme, while reorganizing the existing channels instead of introducing an additional input channel, further improved performance to a precision of 89% and a recall of 91%. Finally, replacing the single-model design with a dual-resolution architecture (version 4), in which a 5 kbp window model detects SVs smaller than 2 kbp and a 50 kbp window model captures larger events that would otherwise be fragmented across multiple images, produced the best results, with a precision of 92% and a recall of 94% (Table 1).

**Table 1.** Impact of input-feature design on SV-calling performance. Successive versions of the pileup-image representation were evaluated on HG002 chromosomes 12–22 at 28× coverage. Adding explicit coverage information, inter-alignment signatures, and dual-resolution models progressively improved performance, with the final ARCLID configuration achieving the highest precision, recall, and F1 score.

| Configuration and Input Features | Precision(%) | Recall(%) | F1(%) |
| --- | --- | --- | --- |
| **V1:** intra-alignment + combination of qs and mp + coverage values in first row | 81 | 83 | 82 |
| **V2:** V1 + coverage line | 85 | 87 | 86 |
| **V3:** V2 + inter-alignment information | 89 | 91 | 90 |
| **V4:** V3 + dual-resolution models (final ARCLID) | 92 | 94 | 93 |

### Resource Usage Evaluation

We benchmarked ARCLID against other SV callers in terms of wall-clock time, CPU time, and memory usage using the HG002 28× across different numbers of cores and threads. The evaluation was performed on a high-performance computing (HPC) system equipped with an NVIDIA V100 GPU, as ARCLID relies on two deep learning models and requires GPU acceleration. Under this setup, ARCLID completed SV calling in approximately 60 minutes with 16 threads. In comparison, Sniffles2 was the fastest SV caller across various computational settings. Overall, ARCLID demonstrates relatively efficient scalability across computational settings and provides a practical solution for researchers. Full evaluation results, including detailed resource usage, are reported in Supplementary Table 21.

## Discussion

In this study, we introduce ARCLID, a structural variant caller built to remain reliable as sequencing coverage decreases. This work also yields a resource of independent value. Reliable benchmarking beyond HG002 is limited by a scarcity of high confidence truth regions in other samples. We curated the HGSVC call sets for HG00733 and NA19240, excluding densely clustered variation and sites with missing or partial genotypes to produce cleaned, high confidence benchmarking regions for both genomes. We release these publicly as a community resource. Together with the HG002 GIAB set, they extend rigorous, low ambiguity structural variant evaluation to two additional, ancestrally distinct genomes and give the community a stronger basis for assessing future callers.

ARCLID extracts informative signals from aligned reads and structures them as two-dimensional pileup images, which are then processed by two deep learning models based on YOLOv11x to identify, classify, and genotype SVs in a single framework. By framing SV detection as an object detection problem, similar to approaches in computer vision, ARCLID learns both low level and high level visual features in the pileup representation and captures variant signatures across a wide range of sizes. Across comparisons with widely used callers, including Sniffles2, cuteSV, SVIM, pbsv, Svision-pro, and DeBreak, these established methods lose sensitivity at lower coverages and for very large variants. ARCLID is most distinctive in this low coverage regime, where it preserves a strong balance of precision and recall as depth falls and other tools decline sharply. At high coverage ARCLID is competitive with the best performing methods rather than uniformly superior, which is consistent with its design focus. ARCLID also localizes breakpoints accurately, which supports downstream analyses such as exon disruption assessment and copy number estimation where precise boundaries matter.

ARCLID’s robustness is most valuable where sequencing data are sparse, such as cost-constrained population genomics studies that trade sequencing depth for population breadth [41]; clinical settings with limited or degraded DNA from small biopsies or FFPE samples [42]; and outbreak investigations requiring rapid, low-cost sequencing to guide public health action [43]. These observations raise a practical question, whether improvements in SV calling algorithms, rather than increases in sequencing coverage, can offer a more cost-effective route to high quality variant discovery.

Sequencing costs are a critical consideration in large-scale and clinical genomics projects. For example, Simonetti *et al.* [44] mentioned that the COVseq protocol for SARS-CoV-2 surveillance multiplexes hundreds of samples on a single run, allocating only ∼0.5–1 million reads per sample (a modest read budget), yet still achieves ≥10× coverage across ≥95 % of the 29 kb genome for under US $15 per genome. Also, according to Genohub [45], whole-genome sequencing of a human sample at ∼30× coverage with the PacBio Revio system, including library preparation, costs approximately US $2,000, producing ∼90 Gb of HiFi data. In comparison, the older Sequel II system would require three SMRT cells to achieve similar throughput, at a cost of around US $8,700 (https://genohub.com/ngs/, accessed August 2025). While ∼ 30× coverage has traditionally been viewed as the benchmark for accurate SV detection, ARCLID’s robustness at lower coverages challenges this paradigm.

ARCLID currently focuses on insertion and deletion detection from PacBio HiFi data, where it maintains strong performance across coverage levels. Our central aim was to demonstrate that framing SV detection as an object detection problem over pileup images is a viable and effective approach. Central to this approach is the way read derived signals are encoded into the pileup representation. ARCLID arranges three complementary signals in pileup images (intra alignment signals, inter alignment signals, and coverage line). We present these design choices not only to explain ARCLID’s performance but to show the community how deliberate feature encoding in pileup images can drive detection accuracy, and to encourage further exploration of which signals matter most. Support for Oxford Nanopore Technologies data and for additional structural variant classes such as inversions and translocations is a future direction of this work.

## Acknowledgements

The authors acknowledge DTU computing center for enabling the use of high-performance computing clusters (HPC) for this research. The authors received no financial support for the research, authorship, and/or publication of this article. The project has been carried out within the *Bio Digitalization and Data Science Group (BDD Group)* at the Department of Biotechnology and Biomedicine, Technical University of Denmark (DTU). S.M. is supported by the SciLifeLab & Wallenberg Data Driven Life Science Program (grant: KAW 2024.0159)

## Author Contributions

Author Contributions Sajad Tavakoli: Conceptualization; methodology; software; validation; benchmarking; formal analysis; data curation and annotation; writing-original draft; writing-review; visualization; project administration. Rasmus J. N. Frandsen: Conceptualization; project administration; supervision; validation; writing-editing; writing-review. Sina Majidian: Validation, writing-editing; writing-review. Marjan Mansourvar: Conceptualization; project administration; supervision; validation; writing-review.

## Data Availability

All data used in this study are publicly available and can be downloaded from the following links:

- Reference Genome hg37 and hg38:

- hg37: ftp://ftp-trace.ncbi.nih.gov/1000genomes/ftp/technical/reference/phase2_reference_assembly_sequence/hs37d5.fa.gz
- hg38: http://hgdownload.soe.ucsc.edu/goldenPath/hg38/bigZips/hg38.fa.gz
- HG002 HiFi aligned reads: ftp://ftp-trace.ncbi.nlm.nih.gov/ReferenceSamples/giab/data/AshkenazimTrio/HG002_NA24385_son/PacBio_CCS_15kb/alignment/HG002.Sequel.15kb.pbmm2.hs37d5.whatshap.haplotag.RTG.10x.trio.bam
- HG002 truth set: https://ftp-trace.ncbi.nlm.nih.gov/giab/ftp/data/AshkenazimTrio/analysis/NIST_SVs_Integration_v0.6/HG002_SVs_Tier1_v0.6.vcf.gz
- HG002 high-confidence regions: https://ftp-trace.ncbi.nlm.nih.gov/giab/ftp/data/AshkenazimTrio/analysis/NIST_SVs_Integration_v0.6/HG002_SVs_Tier1_v0.6.bed
- Challenging medically relevant genes (CMRG): https://ftp-trace.ncbi.nlm.nih.gov/ReferenceSamples/giab/release/AshkenazimTrio/HG002_NA24385_son/CMRG_v1.00/
- HG00733 HiFi reads: https://ftp.1000genomes.ebi.ac.uk/vol1/ftp/data_collections/HGSVC2/working/20190925_PUR_PacBio_HiFi/
- NA19240 HiFi reads: https://ftp.1000genomes.ebi.ac.uk/vol1/ftp/data_collections/HGSVC2/working/20191005_YRI_PacBio_NA19240_HiFi/
- HG00733 and NA19240 truth set: https://ftp.1000genomes.ebi.ac.uk/vol1/ftp/data_collections/HGSVC2/working/20210806_PAV_VCF/
- The low-ambiguity SV regions for HG00733 and NA19240 have been uploaded to Zenodo and can be downloaded through this link: https://doi.org/10.5281/zenodo.21879968

## Code Availability

ARCLID is publicly available on GitHub: https://github.com/sajadtavakoli/ARCLID

## Conflict of Interest Statement

The authors declare no conflicts of interest.

## Notes

### Competing Interest Statement

The authors have declared no competing interest.

### Summary of Updates

Main Changes: 1) SVision-pro and DeBreak were added to the benchmarking analysis. 2) The SV regions of HG00733 and NA19240 were curated, and the benchmarking results were updated accordingly. 3) Additional details were added to the Methods section. 4) All visualizations were updated to include the two newly benchmarked tools.

