## Supplementary Information for "Accurate and Robust Characterization of Structural Variants at Low Coverages with ARCLID"

### S1. Deep Learning Model

The deep learning model used in this study to detect structural variants (SVs) is YOLOv11x, which contains approximately 56 million parameters. We used Ultralytics [1] and pytorch libraries in python to train the deep learning model. YOLOv11x is organized into three main components: the backbone, the neck, and the head. During the forward pass, an input image is processed through a series of convolutional layers and CSP-based backbone blocks [2]. These initial layers extract hierarchical features by applying learned filters that capture edges, textures, and higher-order patterns. The backbone employs cross-stage partial (CSP) connections to reduce computational redundancy while preserving feature richness. In addition, YOLOv11x incorporates attention mechanisms, which improve feature selection and representation compared to earlier versions such as YOLOv8, thereby enhancing the model's ability to focus on relevant regions. As the image traverses the backbone, spatial resolution decreases while channel depth increases, enabling the network to capture more abstract semantic information. Intermediate feature maps are then passed to the neck (e.g., PANet), which fuses multi-scale features through upsampling and concatenation, strengthening the model's capability to localize objects of varying sizes.

In the detection head, YOLO generates predictions by applying anchor-free detection mechanisms. Instead of predefined anchor boxes, the model directly predicts bounding box coordinates (x, y, width, height), objectness scores, and class probabilities at three distinct scales (e.g., 80x80, 40x40, 20x20 grids). Each grid cell's output tensor combines localization (via CIoU-aware regression), confidence (probability of containing an object), and classification scores. The model employs a combination of convolutional layers and activation functions (e.g., SiLU) to refine these predictions. Finally, the outputs from all scales are aggregated, and non-maximum suppression (NMS) is applied post-forward pass to filter overlapping detections, yielding the final set of bounding boxes, classes, and confidence scores. This streamlined process ensures real-time efficiency while maintaining high detection accuracy across diverse object sizes.

YOLO employs three primary cost functions to optimize its object detection performance [3]: box loss (CIoU for bounding box regression, weight=7.5), classification loss (BCE for class prediction, weight=0.5), and distribution focal loss (DFL) for precise bounding box distribution modeling (weight=1.5). These default weights, defined in the hyperparameter configuration, balance the contributions of each component during training, ensuring accurate localization, classification, and confidence estimation. The values are tuned to prioritize bounding box accuracy (highest weight for CIoU) while maintaining a balance between detecting objects (objectness) and refining class predictions.

**Box Regression Loss (CIoU).** The Complete Intersection over Union (CIoU) loss enhances the traditional IoU metric by introducing penalties for both the distance between box centers and discrepancies in aspect ratios [3]. Its formula is shown in below.

$$\mathcal{L}_{CIoU} = 1 - \mathrm{IoU} + \frac{\rho^2(b_{pred}, b_{gt})}{c^2} + \alpha v$$

$\rho$ is the Euclidean distance between box centers, $c$ is the diagonal length of the enclosing box C, $v$ is the penalty for aspect ratio mismatch, and $\alpha$ balances the $v$ and center distance terms.

**DFL.** Designed to refine bounding box localization [3], DFL treats coordinates as discrete distributions and emphasizes probabilities near the target value:

$$\mathcal{L}_{DFL} = -\left((y_{right} - y)\log(p_{left}) + (y - y_{left})\log(p_{right})\right)$$

Where $y$ is the continuous target value (e.g., box coordinate), and $y_{left}$, $y_{right}$ are the nearest discrete bins with probabilities $p_{left}$, $and$ $p_{right}$.

**Classification Loss.** This loss function [3] handles multi-class prediction by computing the binary cross-entropy between ground truth labels and predicted class probabilities:

$$\mathcal{L}_{cls} = -\frac{1}{C}\sum_{c=1}^{C}[y_c \log(p_c) + (1 - y_c)\log(1 - p_c)]$$

Where $C$ is the number of classes, $y_c$ is the binary groundtruth label (0 or 1) for class $c$, and $p_c$ is the model's predicted probability for that class.

**Total Loss.** The combined loss is a weighted sum of the individual components as below:

$$\mathcal{L}_{total} = 7.5\mathcal{L}_{CIoU} + 0.5\mathcal{L}_{cls} + 1.5\mathcal{L}_{DFL}$$

**Training process.** Figure 1. a shows the training process before manual curation, where the loss curves were less stable and the validation metrics fluctuated considerably, indicating that inconsistencies between the pileup images and their ground-truth annotations hindered model convergence. In contrast, Figure 1. b shows the training process after manually curating approximately 50,000 images and retaining only those in which the SV signal was correctly

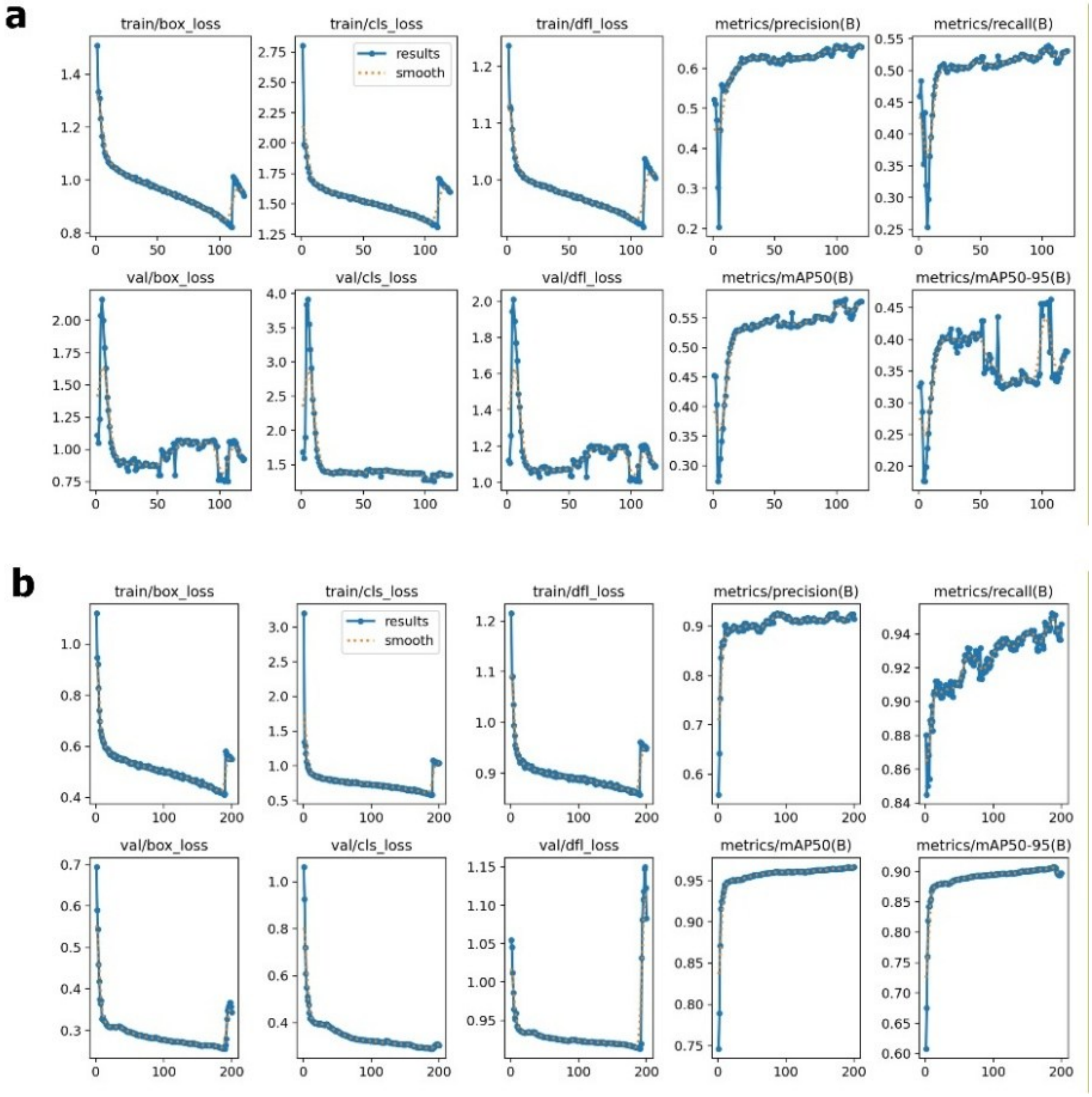

**Supplementary Figure 1. Effect of manual curation on model training. a,** Training before curation. **b,** Training after manual curation of approximately 50,000 pileup images, showing faster and more stable convergence.

aligned with the annotated bounding box. Following curation, the model converged rapidly. Most of the reduction in training and validation losses occurred within the first 10–20 epochs, while precision, recall, and mAP increased sharply during the same early stage. The metrics then improved gradually and remained relatively stable throughout the remaining epochs. These results indicate that manual curation substantially improved label quality, enabling faster, smoother, and more reliable training.

### S2. Execution Parameters

**Truvari v5.0.0**

**Truvari relaxed mode:**

truvari bench -b truth_set.vcf.gz -c pred.vcf.gz -o truvari_results --includebed high_confidence_regions.bed --no-ref b --refdist 1000 --pctseq 0 --passonly

**Truvari strict mode:**

truvari bench -b truth_set.vcf.gz -c pred.vcf.gz -o truvari_results --includebed high_confidence_regions.bed --no-ref b --refdist 50 --pctsize 0.9 --pctovl 0.9 --pctseq 0 --passonly

**Pbmm2:**

pbmm2 align $ref_idx $fq_all $alignment --num-threads 16 --sort --sample $sample_name --preset CCS

**Sniffles2 v2.0.1:**

sniffles --input aligned_reads.bam --vcf output.vcf --genotype-ploidy 2

**cuteSV 1.0.3:**

We set cuteSV parameters along with different min support read given the coverage of the sample using the original paper and GitHub of cuteSV. min_support for different coveages: 28X → 6, 20X → 4, 10X → 3, 5X → 2

cuteSV aligned_reads.bam output.vcf ./ --sample sample_name --max_cluster_bias_INS 1000 --diff_ratio_merging_INS 0.9 --max_cluster_bias_DEL 1000 --diff_ratio_merging_DEL 0.5 --min_support min_sup

**SVIM v2.0.0:**

svim alignment ./ aligned_reads.bam ref.fa --sample HG002 --type DEL,INS --cluster_max_distance 1.4

cat ./variants.vcf | grep -v 'SUPPORT=1;\|SUPPORT=2;\|SUPPORT=3;\|SUPPORT=4;\|SUPPORT=5;\|SUPPORT=6;' > ./variants_filtered.vcf

**PBSV v2.11.0:**

pbsv discover aligned_reads.bam ./vars.svsig.gz --tandem-repeats tnd_file.bed

pbsv call ref.fa ./vars.svsig.gz variants.vcf -t INS,DEL

**Svision-pro v2.5:**

SVision-pro --target_path $bam_file --genome_path $ref --model_path $model_path --out_path $out_path --sample_name $sample --detect_mode germline --preset hifi

**DeBreak v1.0.2:**

debreak --bam ${bam_file} --outpath debreak_out/ --depth ${cov} --rescue_large_ins --rescue_dup --poa --thread 32

**VISOR:**

The entire process of using VISOR to generate simulated reads

1) Rscript /home/user/visor_app/VISOR/scripts/randomregion.r -h
2) cut -f1,2 small.reference.fa.fai > chrom.dim.tsv
3) Rscript /home/user/visor_app/VISOR/scripts/randomregion.r -d chrom.dim.tsv -n 15000 -l 350 -s 50 -x exclude.bed -v 'insertion,deletion,inversion,inverted tandem duplication,translocation copy-paste,reciprocal translocation' -r '40:40:5:5:5:5' | sortBed > HACk.random.bed
4) VISOR HACk -g small.reference.fa -b HACk.random.bed -o hack.3.out
5) VISOR LASeR -g small.reference.fa -s hack.3.out -b shorts.laser.simple.bed -o laser.1.out --threads 4 --coverage 30 --length_mean 13000 --length_stdev 5000 --tag --fastq --compress --read_type pacbio --error_model pacbio2016 --qscore_model pacbio2016
